# Coding odor modality in piriform cortex efficiently with low-dimensional subspaces: a Shared Covariance Decoding approach

**DOI:** 10.1101/2024.06.26.600792

**Authors:** Delaney M. Selb, Andrea K. Barreiro, Shree Hari Gautam, Woodrow L. Shew, Cheng Ly

## Abstract

A fundamental question in neuroscience is how sensory signals are decoded from noisy cortical activity. We address this question in the olfactory system, decoding the route by which odorants arrive into the nasal cavity: through the nostrils (orthonasal), or through the back of the throat (retronasal). We recently showed with modeling and novel experiments on anesthetized rats that orthonasal versus retronasal modality information is encoded in the olfactory bulb (OB, a pre-cortical region). However, key questions remain: is modality information transmitted from OB to anterior piriform cortex (aPC)? How can this information be extracted from a much noisier cortical population with overall less firing? With simultaneous spike recordings of populations of neurons in OB and aPC, we show that an unsupervised and biologically plausible algorithm, Shared Covariance Decoding (SCD), can indeed linearly encode modality in low dimensional subspaces. Specifically, SCD improves encoding of ortho/retro in aPC compared to Fisher’s linear discriminant analysis (LDA). Consistent with our theoretical analysis, when noise correlations between OB and aPC are low and OB well-encodes modality, modality in aPC tends to be encoded optimally with SCD. We observe that with several algorithms (LDA, SCD, optimal) that the decoding accuracy distributions are invariant when GABAA (ant-)agonists (bicuculline and muscimol) are applied to OB, consistent invariance in population firing in aPC. Overall, we show modality information can be encoded efficiently in piriform cortex.

## 1 Introduction

A key question with origins dating back to the beginning of theoretical neuroscience (Barlow, 1961, 1972) is: how are sensory signals extracted from single-trial activity, despite its high variability across repeated trials (Shadlen and Newsome, 1998; Mainen and Sejnowski, 1995; Averbeck et al., 2006; Nolte et al., 2019)? Cortical spikes, which are the fundamental units that carry information, are known to be variable (or ‘noisy’) even at the very beginning layers of cortex. Here we address this question in the olfactory system, assessing how the fidelity of olfactory signals can be maintained as it propagates upstream from a pre-cortical region to a cortical region. Our approach follows a recently hypothesized general decoding scheme, called Shared Covariance Decoding (SCD), which leverages correlations between cortical and pre-cortical neural populations to isolate low-dimensional, low-noise decoding subspaces.

The difficulty of extracting sensory signals is further highlighted by many recent experiments that have shown that early cortical regions have multiple streams of information beyond sensory signals. In primary visual cortex, spiking activity encodes decision making information (Steinmetz et al., 2019; Zhang and Zador, 2023), whisker movement and other motor behaviors (Stringer et al., 2019). In addition to odor identity, olfactory cortex also encodes spatial navigation variables (Poo et al., 2022). Thus, cortical population activity is high-dimensional, containing information from external stimuli mixed with ongoing, recurrent and feedback activity (Arieli et al., 1996). How can sensory information be extracted and made available to higher cortical regions downstream?

In the olfactory system, we specifically consider coding of an important component of olfactory sensory information in anterior piriform cortex (aPC): odor *modality* (as opposed to odor identity, see Barreiro et al. (2024)). By odor modality, we refer to the two mechanisms by which odors can enter the nasal cavity: odors entering from the front of the nasal cavity during sniffing or inhalation are called orthonasal (ortho), while odors entering from the rear during eating and exhaling are called retronasal (retro). Even though olfactory research predominately focuses on ortho stimulation, modality (ortho vs retro) is known to be an important factor in olfaction and flavor perception. Blankenship et al. (2019) showed in transgenic mice that retro stimulation leads to faster association of odors with rewards than with ortho stimulation, as well as modality-dependent activation of other brain regions. Other studies have shown that humans reported different levels of pleasantness of odors depending on modality (Small et al., 2005; Hannum et al., 2021), and that they can distinguish whether an odor was delivered ortho- or retro-nasally without being told the modality (Frasnelli et al., 2008). Since modality is a crucial part of olfactory processing, knowing whether and how it is encoded in aPC would significantly impact our understanding of odor processing.

Although recent work by our labs (Craft et al., 2023, 2021) and others (Scott et al., 2007) have shown that modality information is encoded in the olfactory bulb (OB, a pre-cortical region), this question has not been addressed in olfactory cortex. To this end, we collected spike data from neural populations simultaneously in the OB and aPC in anesthetized rats. While the ethological relevance of ortho versus retro coding is most important in awake animals, here we study anesthetized rats, trading ethological realism for experimental control and precision. This allows us to avoid several potentially confounding factors for coding modality, including the breath cycle and perhaps mechanical top-down signals that control the breath cycle, especially during active sniffing. Our anesthetized recordings allowed us to control for these factors, removing effects of the breath cycle completely by use of a double tracheotomy. We ensured that stimulus timing, air flow speed, air pressure, temperature, and odor concentrations were the same for both ortho and retro stimuli, thus isolating the aspects of modality coding that are due solely to the direction of air flow through the nasal cavity (Fig 1Ai).

**Fig. 1.**
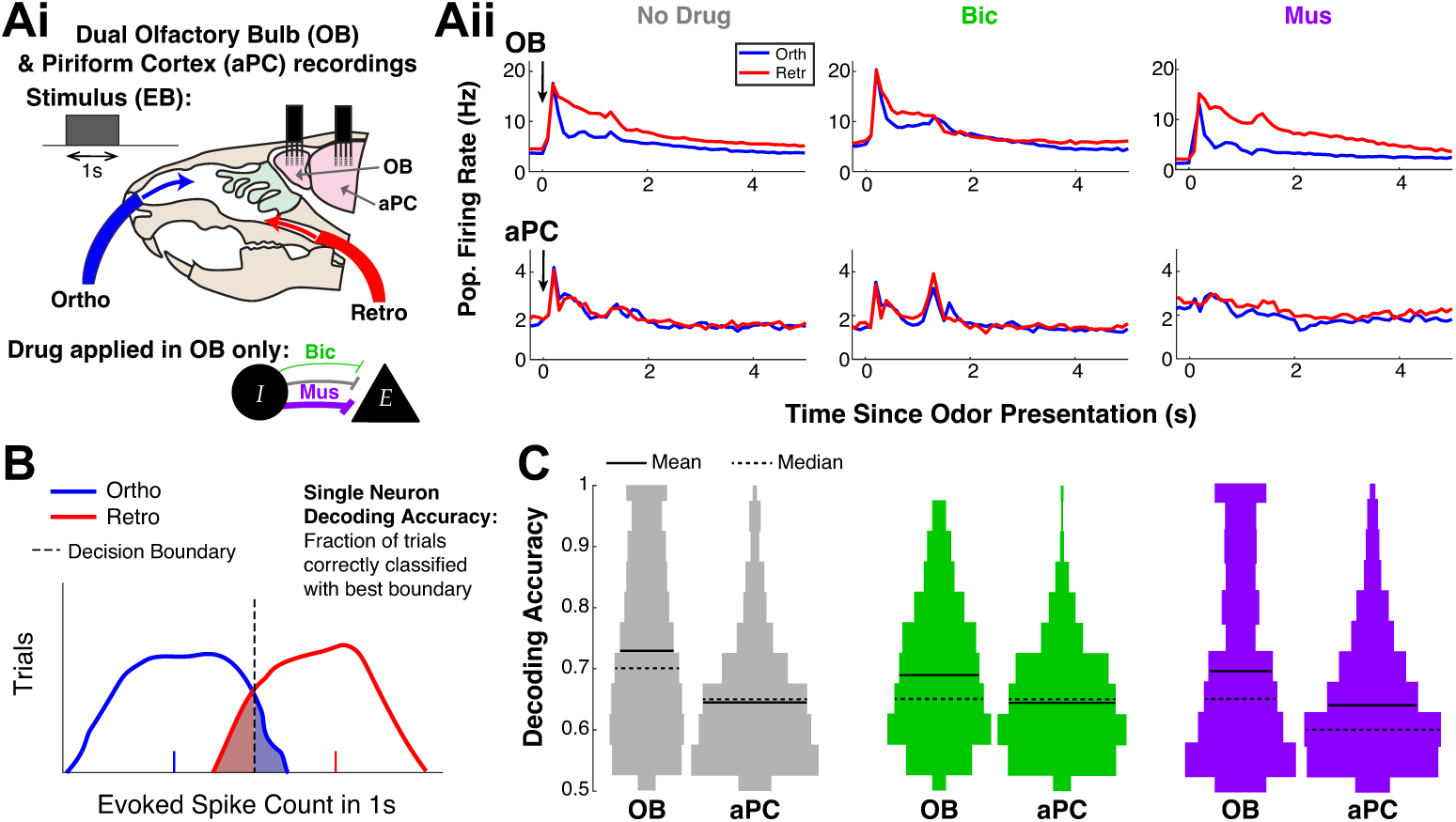
Most single neurons in aPC are poor decoders of ortho versus retro stimuli. Ai) *In-vivo* simultaneous multi-electrode array recordings of anesthetized rats in OB and aPC. Odor stimulation with ethyl butyrate (EB) applied for 1 s with either ortho or retro modality. Drug application in OB only: GABA_A_ antagonist Bicuculline (Bic) diminished inhibition, agonists Musicmol (Mus) enhanced inhibition in OB. Aii) Trial and population-averaged firing rate for each drug preparation shows a clear separation of responses between ortho vs. retro in OB but no separation in aPC. B) Single neuron decoding using the best decision boundary to separate the ortho and retro (evoked) spike count distribution over trials. C) Violin plots of the single neuron decoding accuracy distributions for all recorded neurons (see Table 1 for population size). The single neuron decoding is clearly better in OB than in aPC, on average.

**Table 1.** Number of rats and total number of individual OB and aPC neurons for each drug preparation subject to the food odor ethyl butyrate (EB).

|  | No Drug | Bicuculline | Muscimol |
| --- | --- | --- | --- |
| **Number of Rats** | 8 | 4 | 3 |
| **OB Neurons** | 913 | 413 | 419 |
| **aPC Neurons** | 905 | 496 | 402 |

Our approach is centered on the hypothesis that the cortical high-dimensional noisy neural population activity (Stringer et al., 2019; Manley et al., 2024) can be cleaned up by projecting it onto a subspace with less noise. We recently developed the SCD method to test this hypothesis based on the premise that sensory signals are correlated across pre-cortical and cortical populations (Barreiro et al., 2024). SCD is an unsupervised algorithm that is biophysically plausible (Lipshutz et al., 2021) and based on canonical correlation analysis (Hotelling, 1992). Here we show that modality is well-encoded in cortical subspaces identified by the SCD algorithm. SCD outperforms the standard Fisher’s linear discriminant analysis (LDA), even when GABA_A_ (ant-)agonists drugs are applied in OB. Guided by mathematical calculations, we assess when SCD performs at optimal limits. When noise correlations between OB and aPC are small and OB well-encodes modality, SCD tends to perform close to optimal limits of linear coding. Specifically, we validate theoretical predictions that the noise correlations between OB and aPC are positively correlated with decoding error (defined as a scaled difference between the best decoding accuracy minus the decoding accuracy with SCD). Our results hold in populations of two and three neurons with reasonable decoding accuracy values. Last of all, we observe that the decoding accuracy distributions under several schemes (LDA, SCD, optimal) are invariant with GABA_A_ agonists (muscimol) and antagonists (bicuculline) applied to the OB, which is consistent with invariance of population firing rates in aPC. Our study details how modality information can be encoded efficiently in low-dimensional subspaces in piriform cortex from the ‘bottom-up’ through the olfactory sensory pathway.

## 2 Results

We consider linear decoding of odor modality (ortho vs. retro) by an ideal observer as a way to assess whether this specific component of olfactory sensory information is available to higher cortical brain regions downstream. Across repeated trials, we assume a fixed rule (i.e., threshold, see Fig 1B) for binary classification of modality based on rate coding (London et al., 2010; Shadlen and Newsome, 1998): the total number of spike counts evoked in the 1 s duration of the odor stimulus. Our experiments consist of dual multi-electrode array recordings in both OB and aPC, recording from many neurons simultaneously in both regions (see total number of neurons pooled over all recordings in Table 1). We use a single food-related odor, ethyl butyrate (EB) that smells sweet (like pineapple), because typically only food odors are perceived retronasally (Small et al., 2005), with 20 trials total for each experiment (10 ortho and 10 retro). Odorized air was delivered for 1 s in duration at 1 minute intervals, with a flow rate of 250 ml/min and 1% of saturated vapor. Considering the importance of fast GABA_A_ inhibitory synapses in OB (Tan et al., 2010; Abraham et al., 2010; Schoppa and Westbrook, 1999), we also apply one of two drugs: bicuculline (Bic), a GABA_A_ antagonist that diminishes the effects of fast inhibitory synapses in OB, muscimol (Mus), a GABA_A_ agonist that enhances the effects of fast inhibitory synapses in OB (Fig 1Ai). When we consider pairs and triplets of neurons in our analysis, we only consider such networks that are simultaneously recorded (see Tables 2 and 4).

**Table 2.** Number of simultaneously recorded pairs of neurons.

|  | No Drug | Bicuculline | Muscimol |
| --- | --- | --- | --- |
| **Pairs of Neurons within OB** | 15,328 | 7,705 | 7,480 |
| **Pairs of Neurons within aPC** | 18,440 | 15,803 | 7,590 |
| **1 OB x 1 aPC** | 25,125 | 13,002 | 13,068 |
| **2 OB x 2 aPC, simultaneously recorded** | 4,124,732 | 3,885,574 | 1,987,629 |
| **Viable 2x2 for noise correlation theory** | 1,959 | 1,557 | 1,071 |

**Table 3.** Statistics of (Pearson’s) correlation between error *E* and average noise correlation *ρ*. As predicted by our theory, *E* and *ρ* are undoubtedly positively correlated for all drug preparations. See bottom row in Tables 2 and 4 for network sizes.

| | Correlation($E, \rho$) | $p$-value | 95% Confidence Interval |
| --- | --- | --- | --- |
| No Drug, 2x2 | 0.522 | 0 | (0.489, 0.554) |
| Bicuculline, 2x2 | 0.471 | 0 | (0.431, 0.501) |
| Muscimol, 2x2 | 0.605 | 0 | (0.565, 0.642) |
| No Drug, 3x3 | 0.266 | 0 | (0.211, 0.318) |
| Bicuculline, 3x3 | 0.385 | 0 | (0.332, 0.436) |
| Muscimol, 3x3 | 0.419 | 0 | (0.377, 0.459) |

**Table 4.** Number of simultaneously recorded triplets of neurons.

|  | No Drug | Bicuculline | Muscimol |
| --- | --- | --- | --- |
| Triplets of Neurons within OB | 184,209 | 102,063 | 92,091 |
| Triplets of Neurons within aPC | 265,373 | 343,064 | 100,332 |
| Total 3 OB x 3 aPC, simultaneously recorded | 876,633,480 | 850,900,785 | 466,577,050 |
| 3x3 sub-sample size | 1,349,311 | 1,673,910 | 1,422,287 |
| Viable 3x3 for noise correlation theory | 1,163 | 1,053 | 1,554 |

To get a bird’s eye view of how these two populations respond to directional modality, we first examine the trial- and population-averaged firing rate (Fig 1Aii). The data shows clear differences between OB and aPC. In all drug preparations in OB, the average firing rates between ortho (blue) and retro (red) are well separated and suggest relatively good encoding of modality (on average). In contrast, in aPC ortho and retro firing rates are almost indistinguishable, and overall the firing rates are lower (notice the difference in the scale of the vertical axes). Although it appears as if the aPC population firing rates change with drug application to OB, i.e., different shaped firing rates and peaks in Figure 1Aii, there are no statistically significant differences in either ortho or retro modality with odor stimuli. This is consistent with more extensive analyses on the summed spike counts in 1 s time window (Fig B3) that are the main focus.

As a baseline measure of coding (Rolls and Treves, 2011), we consider the decoding accuracy of modality using a single neuron (Fig 1B) with the best threshold possible. Pooling over the entire population of recorded neurons (see Table 1) yields a distribution of decoding accuracies, illustrated as violin plots in Fig 1C. Clearly, the decoding accuracies are better in OB than in aPC for all three drug preparations (including no drug), which is consistent with the firing rates shown in Fig 1Aii. Since modality is known to be a crucial component of olfactory processing (Blankenship et al., 2019; Small et al., 2005; Hannum et al., 2021; Frasnelli et al., 2008), yet appears to be poorly encoded in single neurons in aPC, we next investigate how modality information could be better encoded in aPC in larger populations/dimensions, i.e., ≥ 2.

We present the Shared Covariance Decoding (SCD) algorithm (Barreiro et al., 2024) as a plausible way for modality to be efficiently coded in low-dimensional sub-spaces. Our method relies on the joint activity of an upstream region (OB) and the region of interest (aPC), with the aPC spike count projected onto a specific hyper-plane (i.e., a line if considering only two-dimensional coding, see Fig 2A, right column); the SCD algorithm uses canonical correlation analysis (CCA) to find hyperplanes (one for OB and one for aPC) that result in the largest possible Pearson’s correla-tion (Hotelling, 1992) between projected aPC activity and projected OB activity. The resulting correlation between the two regions after hyperplane projections is denoted by *R*_1_. Note that recent work by the Chklovskii lab has shown that the projection of neural activity using CCA can be implemented in synapses using STDP (Lipshutz et al., 2021), so SCD is biological plausible. Furthermore, we emphasize that the hyperplane used to project aPC activity does not take into account the ortho/retro classification, so in this sense SCD is an unsupervised algorithm.

**Fig. 2.**
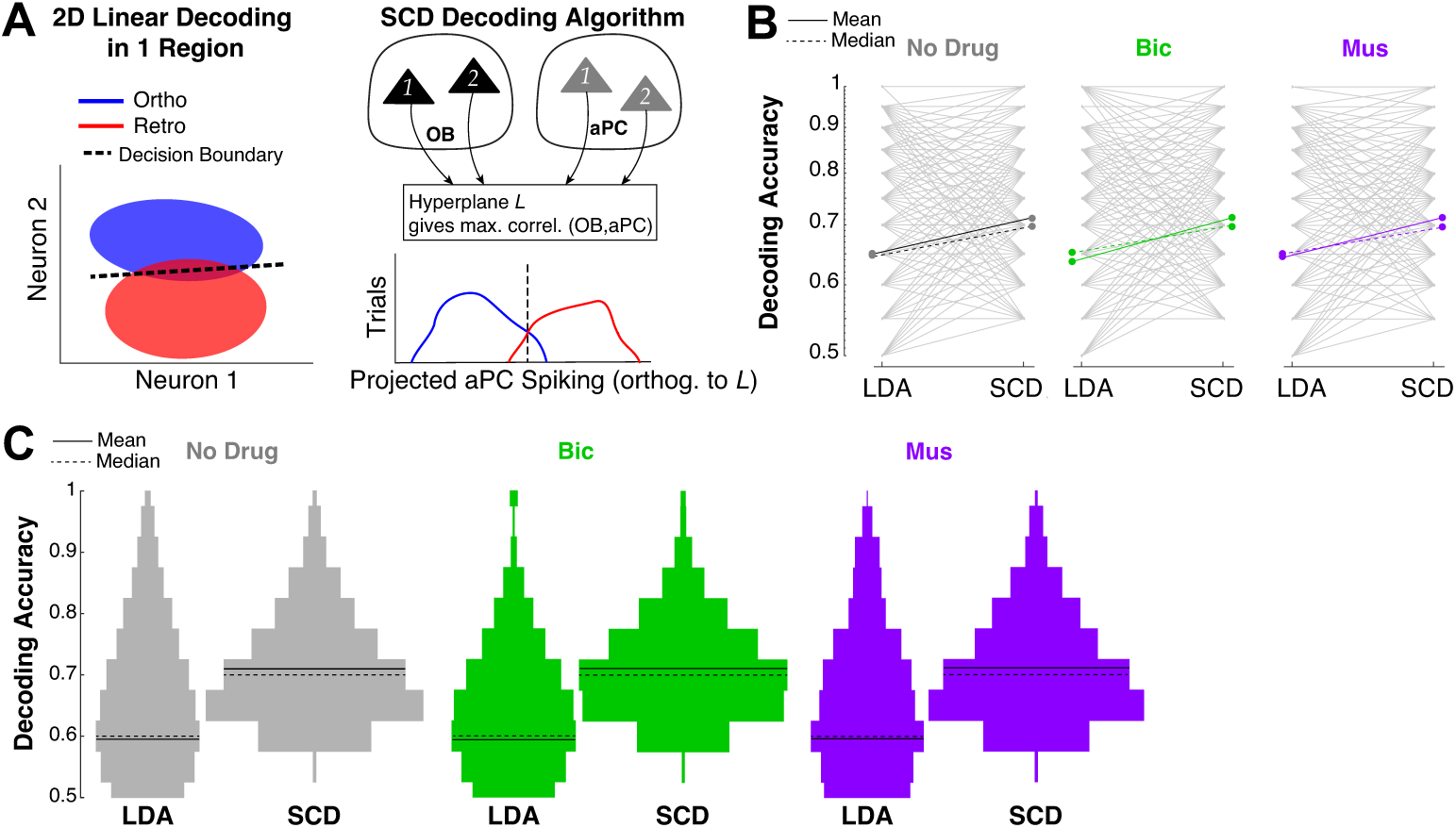
SCD improves aPC decoding of ortho versus retro compared to LDA. A) Left: decoding with a decision boundary as a line (hyperplane) with two neurons, analogous to Fig 1B, using only activity in the specific region. i.e., aPC. Right: idea of the SCD algorithm using two regions: i) use SCD to find hyperplanes *L_OB_*, *L_PC_* such that the OB, aPC activity projected onto *L_OB_* and *L_PC_* respectively is maximally correlated; ii) project aPC spiking activity onto *L_PC_*. B) Comparing SCD with a standard classification algorithm, LDA. SCD can outperform LDA, with robust results across different drug applications in the OB (vertical axis is log-scale). C) Superior performance by SCD compared to LDA is clear in histograms of decoding accuracy over entire 2×2 populations for SCD and all unique pairs of aPC for LDA (histograms shown with violin plots).

To assess how well SCD performs, we compare it to two other linear decoding schemes: i) Fisher’s linear discriminant analysis (LDA), ii) the optimal hyperplane that gives the best possible decoding accuracy. Importantly, both of these schemes are supervised in that they rely on knowing the modality classification of the inputs. LDA is a very common and standard approach which we apply to our data with 10-fold cross-validation. In LDA, the prescribed hyperplane that aPC activity is projected onto is based on the first and second order spike statistics, in particular the mean and variance conditioned on whether the input was ortho/retro are required components (see Methods). LDA happens to be the optimal (best decoding accuracy with a linear decoder) when the spike statistics are normally distributed and assuming homoscedasticity (same covariance matrix with ortho and retro) (Rosenbaum, 2024; Barreiro et al., 2024).

The SCD method performs better than LDA when using two aPC neurons to code modality (Fig 2B, C). In Figure 2B, we show comparison of all possible viable pairs of neurons (two in OB, two in aPC, see fourth row of Table 2); it is evident that on average SCD is better than LDA. Each gray line corresponds to a specific pair of aPC neurons used in both LDA and SCD; note that SCD is coupled to a pair of OB neurons whereas LDA is not. Although at times the gray lines have a negative slope (LDA is better than SCD), the gray lines mostly have a positive slope (SCD is better than LDA). (A minor technical point is that the direct comparison of LDA with SCD in Figure 2B results in ‘over counting’ the number of LDA decoding accuracies because LDA does not depend on the specific OB pair; thus the same line is repeated many time.) Figure 2C demonstrates, with violin plots of the entire (normalized) distribution of decoding accuracies, that SCD is better than LDA when LDA is applied only to the unique pairs of neurons in aPC (see second row of Table 2). Thus, for two-dimensional coding, SCD is overall better than LDA for all three drug preparations. This is remarkable considering that LDA is a supervised method and that it is optimal with Gaussian statistics and homeoscedasticity.

We next surveyed SCD in comparison to the best possible linear decoder. Figure 3A shows, perhaps unsurprisingly, that the optimal linear decoding, calculated by a brute-force algorithm that surveys all possible hyperplanes, is on average much better than SCD. Fortunately, we can address the question of when SCD is (near) optimal with mathematical calculations first presented in Barreiro et al. (2024).

**Fig. 3.**
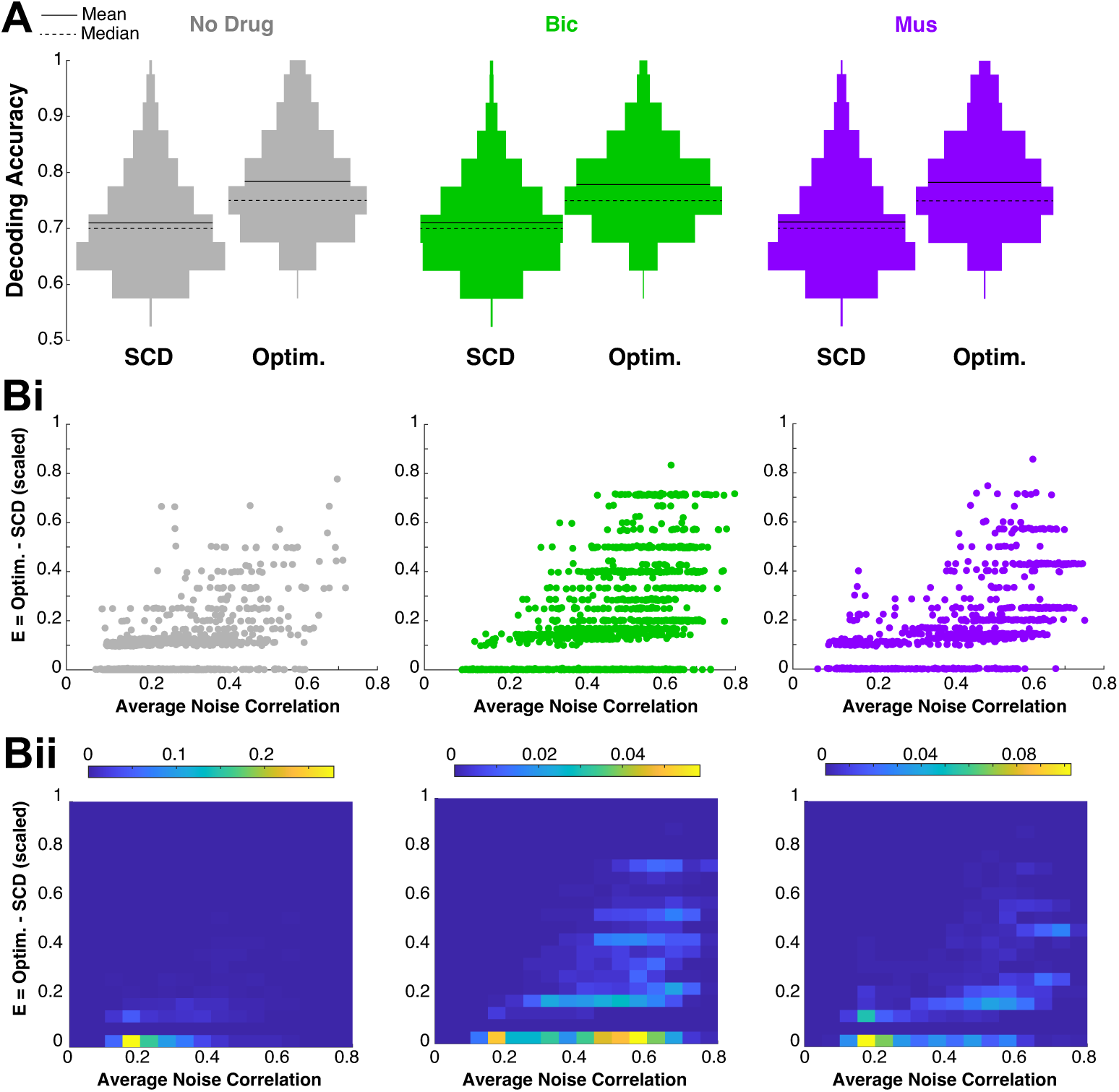
How does SCD compare to optimal decoding for 2×2 pairs? A) Violin plots show, unsurprisingly, that SCD will never outperform the absolute best (optimal decoding). Bi) SCD can perform at the optimal limits (*E* = 0) when the average noise correlations *ρ* are small; *E* [0, 1] is the normalized difference between the optimal decoding and SCD decoding. On the vertical axes is *E* and *ρ* is on horizontal without regard to density of points. As *ρ* increases, *E* tends to increase. Bii) similar to Bi) but shows the actual density of points in a heat map. Overall, *E* is positively correlated with *ρ* (noise correlations between OB and aPC); see Table 3 for detailed statistics.

Assuming Gaussian spike statistics and homeoscedasticity, the SCD method is optimal, i.e., gives largest possible linear decoding accuracy, when the entire noise correlation matrix between the upstream region (OB) and specified region (aPC) is 0.

To apply our theory to this dataset, we must quantify the magnitude of noise correlations between OB and aPC since neural data does not have identically 0 pairwise noise correlations. We also quantify the difference between SCD decoding accuracy and optimal decoding accuracy using a scaled error *E* to measure the difference between SCD and optimal:

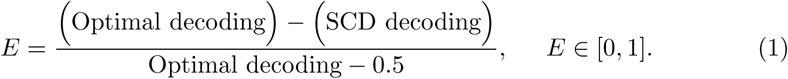

We define the average magnitude of noise correlation *ρ* as follows: let *X_j_* be the *j*^th^ OB neuron’s evoked spike count in the first 1 s after odor onset with for example ortho stimulation and *Y_k_* be the *k*^th^ aPC neuron’s ortho spike count. The noise correlation between this OB and aPC neuron is:

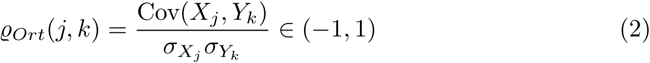

where the Cov and standard deviation *σ* are calculated over trials. For a given 2×2 network (2 OB neurons, 2 aPC neurons), there are 8 total noise correlation values (4 unique pairs across OB and aPC, in each of two modalities: ortho/retro), thus we define:

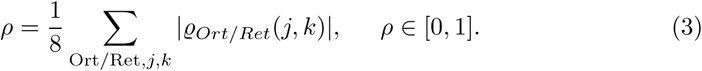

Finally, to isolate the effects that lead to different *E* values, we apply reasonable filters to the 2×2 networks to demonstrate the effectiveness of our mathematical theory. i) The upstream OB region needs to well-encode ortho vs. retro, so we only consider networks where the optimal OB decoding accuracy of the 2×2 network is ≥ 0.75. This is a pragmatic consideration because if there is weak or no signal from OB, then we do not expect information to be accurately decoded downstream using SCD to correlate aPC with OB activity. ii) The resulting projected correlation *R*_1_ from CCA cannot be too small otherwise the high fidelity encoding in OB is diminished, so we only consider networks where *R*_1_ ≥ 0.75. iii) Since we want to use a single scalar to quantify the magnitude of noise correlations *ρ* and there are 8 different values for 2×2 networks, we consider only networks where the standard deviation across all 8 *̺* values is *<* 0.2. The number of viable networks with all of these conditions is much smaller but still large enough for meaningful statistics (see fifth row of Table 2).

With these appropriate conditions applied to the data, our theory predicts that the error and noise correlation will co-vary together. Indeed, Figure 3B shows that average noise correlation magnitude *ρ* (x-axis) and error *E* (y-axis) are positively correlated; Figure Bi shows a scatter plot with a point for each network, and Figure Bii shows the density of points. The summary statistics of this effect in Table 3 (first 3 rows) clearly show positive correlation between *E* and *ρ*. The correlation values are at least 0.471, the statistical significance is 0 (i.e., the probability of the null hypothesis that there is no correlation is 0), and the 95% confidence interval has a tight range.

We apply the same data analysis to 3×3 networks (three OB neurons and three aPC neurons; see Table 4 for counts) and find that our results hold qualitatively but are not as strong. Here the total number of 3×3 networks is prohibitively large due to combinatorial blow-up, so we apply our analysis to a random yet large subset of 3×3 networks (fourth row of Table 4). Similar to 2×2 networks, in 3×3 networks the SCD outperforms LDA, and the optimal outperforms SCD (see Fig 4A). Applying the same filters^1^ as before, it is apparent that the error *E* is positively correlated with average noise correlation magnitude *ρ* (Fig 4B), as in the 2×2 networks. The last 3 rows of Table 3 certainly have positive values but the values are not as large as in the 2×2 (first 3 rows). Appendix A shows how the error *E* is negatively correlated with the CCA-projected correlation *R*_1_ in the 2×2 and 3×3 networks, which is a related prediction based on our theory.

**Fig. 4.**
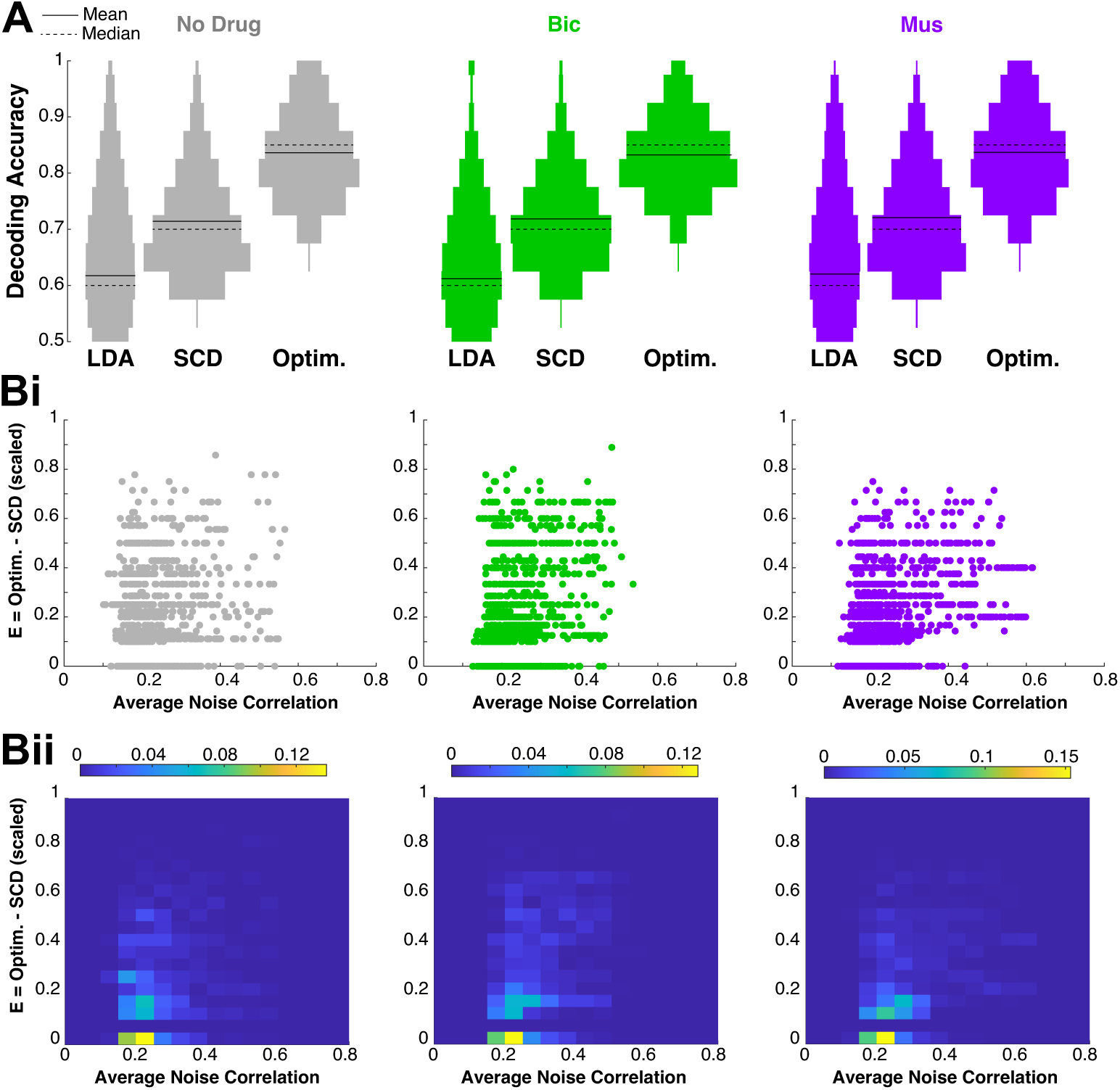
Results for triplets 3×3. The analogous results for more neurons: three in OB and three in aPC. A) The SCD algorithm is much better than LDA, and the optimal is naturally better than SCD. A) SCD can approach optimal limits (*E* = 0) when the average noise correlations *ρ* are small. Both Bi) and Bii) show that *E* is positively correlated with *ρ*; see Table 3 for detailed statistics. There are some differences compared to Fig 3 with the results being not as strong for 3×3 networks, but they generally hold.

Within a given decoding scheme, there are no visual differences across drug preparations for both the 2×2 and 3×3 networks (see histograms of decoding accuracy distributions in Fig 5A,B). The invariance of decoding accuracy in aPC across LDA, SCD and optimal is initially a bit surprising considering how much the two GABA_A_ drugs affect the spiking activity in OB (Craft et al., 2023) and that OB is strongly and recurrently coupled to aPC (Oswald and Urban, 2012; Markopoulos et al., 2012; Boyd et al., 2012). Statistical analyses verify these observations: we use three tests (*t* test assuming unequal variance, Wilcoxon Rank Sum Test, one-way ANOVA) to assess whether the sample means of decoding accuracy distributions are different for each algorithm and find, despite some small *p* values, the effect sizes are beyond minuscule on the order of 0.01 (see Methods; values of up to 0.2 are still considered small in *t* test and Wilcoxon Rank Sum, for one-way ANOVA the effect sizes are orders of magnitude smaller than 0.01), with the following exceptions: LDA for pairs (different means for ND and Bic significant only with *t* test, but effect size of 0.16), LDA for triplets (different means for ND and Bic significant only with *t* test but effect size of 0.16, different means for Mus and Bic significant with *t* test but effect size of 0.14). Given the sample sizes, these are relatively small effect sizes. There are 54 total number of pairwise comparisons (3 drug comparisons, 3 algorithms, 3 tests, 2 networks (pairs, triplets)).

**Fig. 5.**
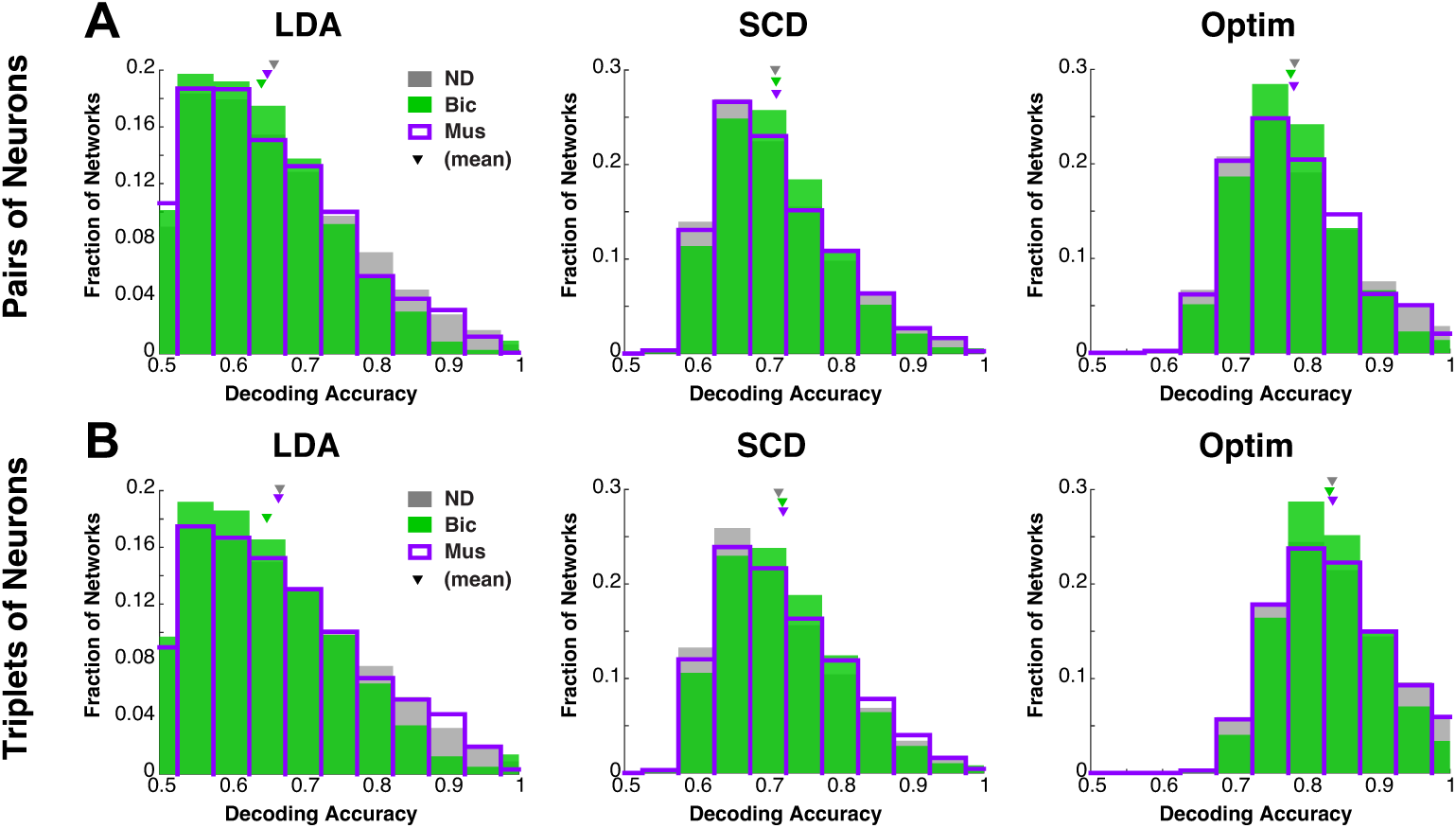
Decoding accuracy is invariant with OB drug application. A) Histograms of decoding accuracies for pairs are very similar with drug application within a given algorithm. B) Same as A) but for triplets.

To better understand why the decoding accuracy distributions are invariant to drug manipulations, we take a closer look at the underlying spike statistics. We do a comprehensive assessment of potential differences with drug preparation and modality, once again finding that besides a few exceptions there are only minuscule effect sizes on the order of 0.01 despite some small *p* values. This assessment includes the mean, variance, covariance, correlation (pooled for all possible neurons/pairs within aPC), as well as the cross-covariance and cross-correlation (between OB and aPC), testing for: i) differences between drug preparations (ND, Bic, Mus), segmented by modality, ii) differences between ortho versus retro, segmented by drug preparation, using all 3 statistical tests. Despite the larger number of comparisons (162 total, 54 for each statistical test), there were only 3 cases where the *p* value was less than 0.01 and the effect size is not minuscule. They are: aPC correlation with ortho stimulation comparing ND to Bic and Mus to Bic (*p <* 0.01 and effect size is small, *<* 0.2 with *t* test), and aPC correlation with retro stimulation comparing ND to Bic (see Methods). See Appendix B and Figure B3 for details. This is in contrast to spike statistics in OB where we have previously noted that GABA_A_ has significant effects (Craft et al., 2023). The invariance of the underlying spike count statistics across drug preps is consistent with the invariance of decoding accuracy distributions.

The average decoding accuracy does not change much in SCD as network size varies from 2×2 (Fig 3) to 3×3 (Fig 4), but this also is true for LDA and optimal, suggesting that this observation might generalize beyond these coding schemes. We speculate that this stems from having different neurons that have a wide range of modality encoding (from bad to good). Intuitively, larger networks should be able to take advantage of more information, at least with the perfect (nonlinear) coding scheme. But perhaps this is not surprising when considering that odor modality is just one aspect of odor information, and there is much more these neurons are encoding beyond modality.

The sparseness of aPC coding has been demonstrated by several labs (Poo and Isaacson, 2009; Isaacson, 2010; Stettler and Axel, 2009; Miura et al., 2012; Otazu et al., 2015), although see Bolding and Franks (2017). There are different measures of sparsity (Willmore and Tolhurst, 2001; Kloppenburg and Nawrot, 2014), ranging from the population kurtosis where higher values correspond to many smaller events, i.e., more sparse, to Treves-Rolles’ formula (Treves and Rolls, 1991). Here we simply use the percent of neurons that spike in a given trial out of the total number recorded, which has previously been reported to range from 3% to 15% with *in vivo* 2-photon calcium imaging (Isaacson, 2010). In comparison, our recordings generally have 10% of aPC spiking to upwards of 40%, with more neurons responding with retro than ortho (Fig B4). So our recordings exhibit denser coding of odors than most of these studies, which is at least consistent with Bolding and Franks (2017).

Although in our controlled experiments the odor is applied for 1 s, the odor duration in awake behaving (sniffing) mammals is on a shorter time scale of hundreds of milliseconds. We applied all 3 decoding schemes (LDA, SCD, optimal) to spike counts in varying window sizes of 100 ms to 1 s in increments of 100 ms to determine whether the results are significantly different. We use the same 2 cell networks consisting of all possible pairs and triplets randomly sub-sampled with the same procedure as before. Fig 6 shows that our results hold over many time windows; the average decoding accuracies have consistent ordering: LDA *<* SCD *<* optimal, the standard deviation of the decoding accuracies across all networks does not deviate much with time window, and that longer time windows result in a slight increase of decoding accuracy on average (the exceptions are: LDA with Bic, SCD with Mus). Thus, the time window we mainly focus on (1 s) that corresponds to the duration of the odor stimuli in our experiments is the most appropriate, and generally has the highest average decoding accuracy.

**Fig. 6.**
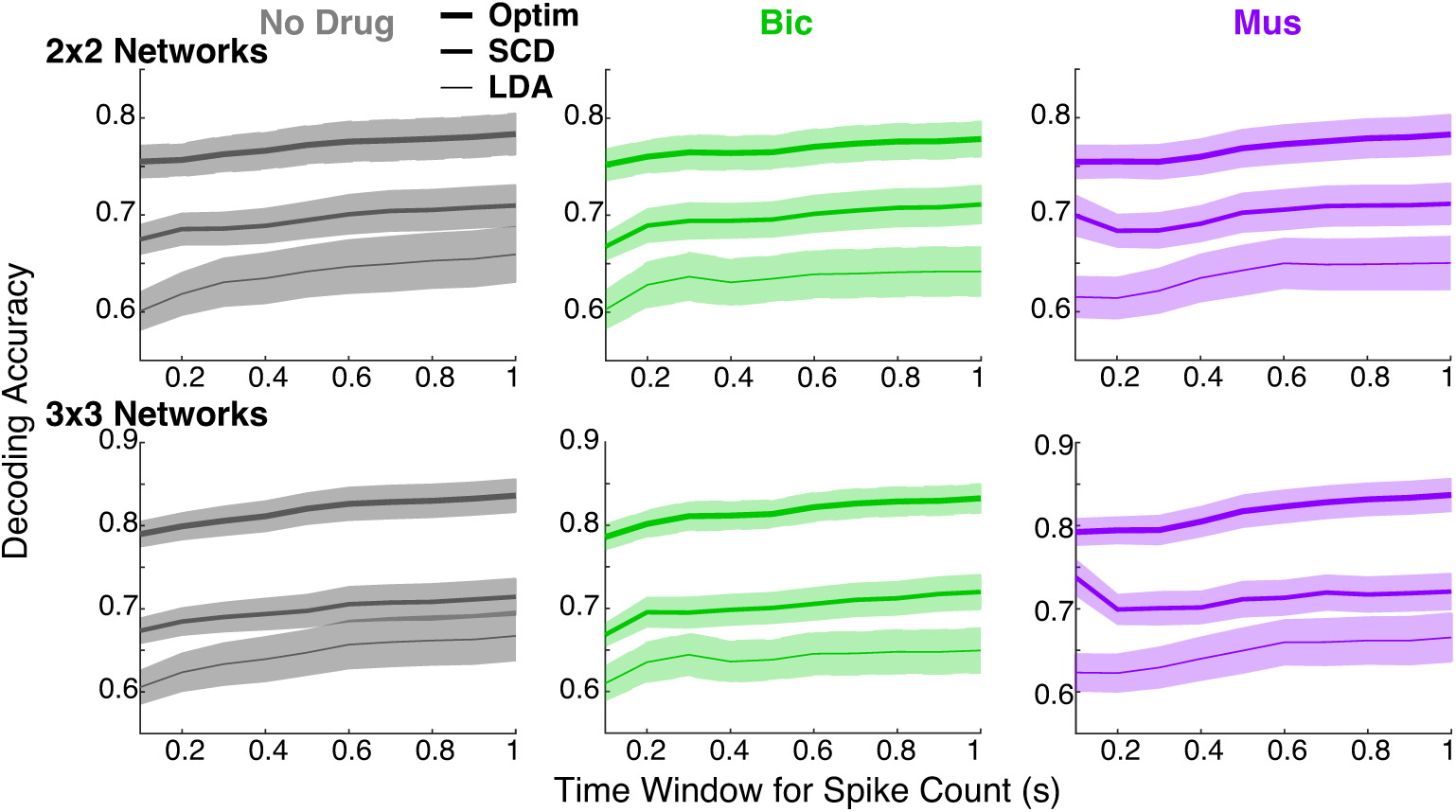
Decoding accuracies with different time bins. Summary of decoding accuracies for summed spike counts in different time windows after odor onset, varying from 100 ms to 1 s in 100 ms increments. Top row shows the average decoding accuracy (lines) among all possible pairs in different drug preps, the shaded region is ±0.25 std dev. Bottom row shows analogous results for triplets. In all cases, the SCD is statistically significantly better than LDA, and optimal is better than SCD; *p*−values are all numerically 0.

## 3 Discussion

Using dual multi-electrode array recordings on tracheotomized anesthetized rats to remove confounds of breathing, combined with computational methods, we show that low-dimensional subspaces can linearly encode modality with decoding accuracy values that vary from reasonable to very accurate. We specifically show that our recently developed SCD algorithm (Barreiro et al., 2024) performs better than LDA across many different time windows, and also verify trends predicted by mathematical theory for when the SCD algorithm performs optimally. The distribution of decoding accuracies with 3 algorithms (LDA, SCD, optimal linear) are invariant with GABA_A_ synaptic agonist and antagonist in precortical OB region over several different networks. The SCD decoding algorithm has now proven to be robust with successful applications to orientation tuning in the visual system, odor identity (Barreiro et al., 2024), and now odor modality.

Considering the success of the SCD algorithm, it is worth taking a closer look at how real neural systems might implement this coding strategy. First, SCD is fully unsupervised; it is based solely on correlations between cortical neurons and the upstream neurons that provide input to cortex. The SCD subspace is defined without any knowledge of the stimulus types (ortho versus retro olfactory cues here) that ultimately are decoded. Second, several theoretical studies have shown that realistic activity-dependent plasticity rules can generate synaptic weights that implement canonical correlation analysis (Gou and Fyfe, 2004; Pehlevan et al., 2020; Lipshutz et al., 2021). In this theory, a network of excitatory and inhibitory neurons (like those we measured here from aPC) that receives input from another population (like the OB neurons we measured here) can perform CCA. This neural algorithm does not require a memory of previous observations to be stored and is performed“on-the-fly”. The synaptic plasticity rules are based on locally-available information about the pre- and postsynaptic neurons activity. This theory suggest that biophysically-realistic implementation of SCD is plausible.

Although the SCD algorithm has been reported in a prior publication (Barreiro et al., 2024), there are numerous differences between that paper and this one. Barreiro et al. (2024) applied the SCD algorithm to awake mice and considered binary classification of basic sensory stimuli: decoding orientation angle in primary visual cortex and odor identity in piriform cortex. Note that these important sensory signals are well-known and heavily studied, whereas odor modality as part of odor input is understudied despite the significant implications it may have on downstream cortical processing. To our knowledge, this is the first study to demonstrate that olfactory cortex codes modality information from the ‘bottom-up’ through the olfactory sensory pathway.

We apply the SCD algorithm to anesthetized rats for reasons previously outlined, but a well-known phenomena with urethane anesthesia is the larger fluctuations, i.e., noise correlations, across trials in neural activity. Despite the larger magnitudes of noise correlations observed here (in comparison to awake data used in Barreiro et al. (2024)), the theory for when SCD is optimal generally holds. Thus, this study enlarges the sphere of experimental recordings and settings in which SCD has been shown to be a viable strategy.

We previously showed, using multi-electrode array recordings of OB in anesthetized rats that modality is encoded relatively accurately even with single neurons, and that a GABA_A_ agonist and antagonist slightly altered the fidelity of ortho/retro information. We found that intermediate levels of inhibition (i.e., no drug) gave the best average decoding accuracy of ortho vs retro modality in OB, and explained our results with a firing rate model that captured many details of our experimental data (Craft et al., 2023). The results in this paper shed light on the details of modality information coding downstream in aPC. Although optimal decoding with single neurons in aPC did not perform well (Fig 1C), simply adding one or two dimensions (neurons) vastly improved coding odor modality with SCD (Fig 2C, 3A). We only consider networks of two or three neurons (four and six for SCD if counting OB neurons) because this suffices to demonstrate our point that odor modality is well-encoded in low-dimensional subspaces. Given the paucity of trials with only ten for each classification, considering larger-sized networks might yield unreasonably good results to this binary classification problem. Since our recordings certainly contain only a subset of neurons, there are undoubtedly many other unrecorded neurons that could implement SCD for odor modality and other sensory signals so that the (relatively) small network sizes should not be viewed as a limitation. One avenue for future work is to extend SCD to a temporally-dynamic projection subspace by using time-dependent canonical correlation analysis (Cao et al., 2019); however, this would require estimation of an increased number of parameters and thus suffer the same estimation issues as a larger (e.g. 4x4) network.

A key feature of odor stimuli is that corresponding OB activity depends on odor concentration (Tan et al, 2010), while there is evidence that coding in aPC might be invariant to concentration (Bolding and Franks, 2018). Here we have shown that with the SCD method for 1 concentration (1% saturated vapor) and 1 odor (EB) that it is possible in principle to use both OB and aPC effectively in an unsupervised manner. The robustness of our results with many concentrations and different odors was not fully explored here. Also even in a single trial, the temporal dynamics of neural activity varies. Extending the SCD method to properly account for time-varying (and concentration varying) inputs, beyond summing spike counts in various time windows (Fig 6), as mentioned above say with time-varying CCA, could be a valuable future contribution.

The invariance of decoding accuracy distributions (Fig 5A,B) is initially a bit surprising since, even though the drug application is in OB, these two regions are tightly recurrently coupled (Boyd et al., 2012; Markopoulos et al., 2012; Oswald and Urban, 2012). But a closer look at the statistics of aPC with a focus on the differences between drug preparations, or differences in modality within a given drug preparation, revealed very few differences among the pooled populations of aPC neurons, and the cross-covariance/correlation of spiking between OB and aPC. Other aspects of odor information such as concentration are known to manifest differently in the OB versus olfactory cortex (Bolding and Franks, 2018). The relatively invariant spike statistics across drug preparations are consistent with invariance of coding performance, thus we did not consider detailed biophysical modeling of the olfactory system (Li and Cleland, 2013, 2017; Craft and Ly, 2022; Craft et al., 2021) to relate spike statistics to decoding performance (Craft et al., 2023). This could be viewed as a limitation: we did not fully explain the underlying mechanistic reasons for how a large change in OB spike statistics could result in little change in aPC spiking. This issue might be addressed in a future study with a mean-field analysis of 2 reciprocally coupled populations (Barreiro et al., 2017; Craft et al., 2021; Schwalger and Chizhov, 2019; Ly and Tranchina, 2007), but is beyond the scope of this current study. Cang and Isaacson (2003) previously reported that granule cells in OB are less active in anesthetized rats than in awake rats, which would be a key factory to consider in future studies.

## Methods

### Ethics statement

All procedures were carried out in accordance with the recommendations in the Guide for the Care and Use of Laboratory Animals of the National Institutes of Health and approved by University of Arkansas Institutional Animal Care and Use Committee (protocol #14049). Isoflurane and urethane anesthesia were used and urethane overdose was used for euthanasia.

### Data and code availability

See https://github.com/chengly70/SCDmodality for MATLAB code implementing all computational components in this paper.

See https://doi.org/10.6084/m9.figshare.14877780.v1 for raw data collected from the Shew Lab.

### Electrophysiological recordings

Data were collected from 8 adult male rats (240-427 g; *Rattus Norvegicus*, Sprague-Dawley outbred, Harlan Laboratories, TX, USA) housed in an environment of controlled humidity (60%) and temperature (23^◦^C) with 12h light-dark cycles. The experiments were performed in the light phase.

#### Surgical preparations

Anesthesia was induced with isoflurane inhalation and maintained with urethane (1.5 g/kg body weight (bw) dissolved in saline, intraperitoneal injection (ip)). Dexamethasone (2 mg/kg bw, ip) and atropine sulphate (0.4 mg/kg bw, ip) were administered before performing surgical procedures. Throughout surgery and electrophysiological recordings, core body temperature was maintained at 37^◦^C with a thermostatically controlled heating pad. To isolate the effects of olfactory stimulation from breath-related effects, we performed a double tracheotomy surgery as described previously Gautam and Verhagen (2012). A Teflon tube (OD 2.1 mm, upper tracheotomy tube) was inserted 10 mm into the nasopharynx through the rostral end of the tracheal cut. Another Teflon tube (OD 2.3 mm, lower tracheotomy tube) was inserted into the caudal end of the tracheal cut to allow breathing, with the breath bypassing the nasal cavity. Both tubes were fixed and sealed to the tissues using surgical thread. Local anesthetic (2% Lidocaine) was applied at all pressure points and incisions. Subsequently, two craniotomies were performed on the dorsal surface of the skull. One craniotomy was over the right olfactory bulb (2 mm × 2 mm, centered 8.5 mm rostral to bregma and 1.5 mm lateral from midline). The other 2 mm × 2 mm craniotomy, for the aPC probe, was positioned 1.5 mm caudal to Bregma, 5.5 mm lateral to the midline.

#### Olfactory stimulation

A Teflon tube was inserted into the right nostril to deliver orthonasal stimuli, and the left nostril was sealed by suturing. The upper tracheotomy tube inserted into the nasopharynx was used to deliver odor stimuli retronasally. Odorized air was delivered for 1 second in duration at 1 minute intervals, with a flow rate of 250 ml/min and 1% of saturated vapor. Two odors were used: ethyl butyrate (EB, a food odor) and 1-Hexanol (Hex, a non-food odor). For a given odor, we used two routes of stimulation, orthonasally and retronasally, for a total of 20 trials, 10 for each ortho and retro, with the order alternating (10 ortho before 10 retro, then 10 retro before 10 ortho trials).

#### Electrophysiology

A 32-channel microelectrode array (MEA, A4×2tet, NeuroNexus, MI, USA) was inserted 400 *µ*m deep from dorsal surface of OB targeting tufted and mitral cell populations. The MEA probe consisted of 4 shanks (thickness: 15 *µ*m, inter-shank spacing: 200 *µ*m), each with eight iridium recording sites arranged in two tetrode groups near the shank tip (inter-tetrode spacing: 150 *µ*m, within tetrode spacing 25 *µ*m). A second 32-channel array (Buzsaki32L, Neuronexus, MI, USA) was inserted through the dorsal surface of the brain to a depth of 6.8 mm stereotaxically targeting anterior piriform cortex. The MEA had 4 shanks (thickness: 50 *µ*m, inter-shank spacing: 200 *µ*m), each with eight iridium recording sites spanning 120 *µ*m at the shank tip. Voltage was measured with respect to an AgCl ground pellet placed in the saline-soaked gel foams covering the exposed brain surface around the inserted MEAs. Voltages were digitized with 30 kHz sample rate (Cereplex + Cerebus, Blackrock Microsystems, UT, USA). Recordings were band-pass filtered between 300 and 3000 Hz and semiautomatic spike sorting was performed using Klustakwik software, which is well suited to the type of electrode arrays used here Rossant et al. (2016).

#### Pharmacology

We topically applied GABA antagonists (bicuculline 20 *µ*M) or agonists (muscimol 20 *µ*M) dissolved in artificial cerebrospinal fluid (ACSF). Gel foam pieces soaked in the ACSF+drug solutions were placed on the surface of OB surrounding the recording electrodes. Note that the concentration of (ant-)agonists can vary over orders of magnitude; some labs use 10 *µ*M (Li et al., 2019; Sharp and Finger, 2002) and others use up to 10 to 10^4^ higher concentration levels (Wachowiak and Cohen, 1999; Kollo et al., 2014). The gel foams were placed 10 minutes before beginning a recording.

### Data analysis

Data was collected *in vivo* from the mitral cell (MC) layer in the olfactory bulb (OB) and in the anterior piriform cortex (aPC) from multiple anesthetised rats using dual multi-electrode array recording. The data we analyzed consisted of spike recordings of both OB and aPC subject to only EB odor because food odors are perceived ortho- and retronasally (Small et al., 2005); non-food odors do not naturally occur retronasally. The spike counts were calculated using 100 ms overlapping time windows. Three separate drug preparations were used in order to analyze inhibitory effects on MC spiking responses: no drug (control), Bicuculline (GABA_A_ antagonist, i.e., decreasing inhibition), and Muscimol (GABA_A_ agonist, i.e., increasing inhibition). Data was obtained from 8 viable rats in total. After spike sorting, we identified and eliminated units (cells) that had firing rates below 0.008 Hz or more than 49 Hz, calculated over the entire recording duration. We took this conservative approach to eliminate potential artifacts yielding units that had unrealistically high or low firing; see Table 1 for the reported number of total rats and individual cells for each drug preparation. Note that some cells around the odor onset (*t* = 0) will have much larger (*>* 49 Hz) or smaller firing rates than our criteria to eliminate units, but the entire recording duration is much longer with approximately a minute between trials. This is the same procedure we have consistently used in prior publications that contained part of this data (Barreiro et al., 2017; Craft et al., 2021, 2023).

The number of neurons recorded per rat, and the number of recording sessions in a rat varied by region (OB vs aPC) and drug type. For ND, among the 8 total rats with anywhere from 3 to 5 recordings, the mean±std per rat were 114.13±22.91 neurons in OB and 113.13±78.19 neurons in aPC recorded. For Bic, among the 4 total rats with 2 to 4 recordings per rat, the mean std were 103.25 30.45 neurons in OB and 124.00±132.33 neurons in aPC recorded. For Mus, among the 3 total rats with 3 or 6 recordings per rat, the mean std were 139.67 -47.50 neurons in OB and 134.00±120.83 neurons in aPC recorded. More details can be found on the dedicated GitHub page with the variables dSizeCells perRecord.mat (for OB) and dSize PC perRecord.mat.

When considering pairs of neurons (2 from OB, 2 from aPC), we looped through all possible pairs but chose to skip over the 2×2 networks where either the OB or aPC spike count matrix was not full-rank. The values stated in the fourth row of Table 2 are the final number of 2×2 networks we used. We used the same procedure for 3×3 networks.

### Decoding algorithms

#### Fisher’s Linear Discriminant Analysis (LDA)

LDA for binary classification is well-known; we state the details here for completeness.

Assume *n* neurons in a given network with 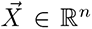 denoting a random variable of spike counts in the 1 s odor-evoked period, and 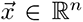 be a particular observation of spike counts in a given trial. Let the conditioned mean of spike counts (taken over trials) be 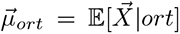 and 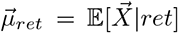, and similarly the conditioned co-variance matrix for ortho stimulation is denoted by Σ*_ort_* with the (*j, k*) entry defined as: 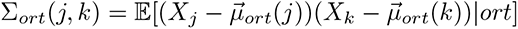 (similarly for retro). Let

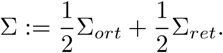

The algorithm projects a given observation 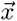 onto the vector

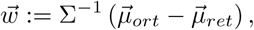

classifying observation 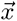 as ortho if:

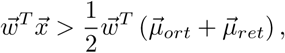

and classifying the observation 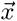 as retro if

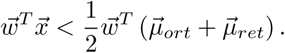

This is a standard criteria derived by the Bayesian criteria 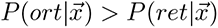 for say ortho classification, assuming *P* (*ort*) = *P* (*ret*), which we have since there are ten trials of each modality. This is implemented in MATLAB with the functions `fitcdiscr` and `kfoldPredict` with 10-fold cross-validation.

#### Shared Covariance Decoding (SCD)

Let *X* and *Y* be matrix of spike counts in OB and aPC, respectively, where the (*j, k*) entry of *X* corresponds to the *j*^th^ trial of the *k*^th^ OB neuron’s spike count in the 1 s odor-evoked period (similarly for *Y*). Both matrices are of size 20×*n* assuming 20 total trials and *n* neurons in each region. The CCA algorithm (Hotelling, 1992) gives the vectors 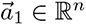 and 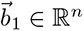 (commonly called the ‘Sample Canonical Coefficients’ of *X* (resp. *Y*)) that maximize the Pearson’s correlation between 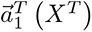 and  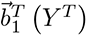, which are vectors of size 20 × 1. We label this maximum Pearson’s correlation by *R*_1_; there are subsequent vectors 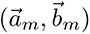 that give projected correlations *R_m_ < R*_1_, but are not used in the SCD algorithm.

Since we are only focused on aPC decoding, we consider the hyperplane projected spike counts:  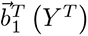 (commonly called ‘Canonical Scores’ of *Y*) and calculate the decoding accuracy with the best threshold. We emphasize that the vector 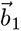 is calculated in an unsupervised manner without knowing the ortho/retro labels of the individual trials.

We use the MATLAB function canoncorr which conveniently outputs the largest canonical correlation *R*_1_ and the Canonical scores 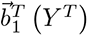.

#### Statistical significance and effect size

We use three tests to assess whether a given effect is statistically significant with different population averages: 1) two-sample *t* test with unequal variance, 2) Wilcoxon rank sum test, 3) one-way ANOVA. Each of these tests are used to rule out the null hypothesis that the population means are the same in two different categories (Cohen, 2013). Each test is different and has various underlying assumptions: 1) normal samples, 2) non-parametric test assuming independent groups and equal variance, 3) including sample variance to assess differences in means assuming equal variance and normally distributed residuals. We consider three tests, as opposed to one, for robustness.

In addition to the *p* values that indicate the probability of the null hypothesis holding (no effect, same population means), the ‘Effect Size’ is an important consideration. We use the conventions outlined in Cohen (2013) and Tomczak and Tomczak (2014) that provide formulas for effect sizes, and a qualitative ‘rule of thumb’ for when effect sizes are: small, medium or large. For 2 groups of length *n*_1_ and *n*_2_ respectively, the effect sizes we use are:

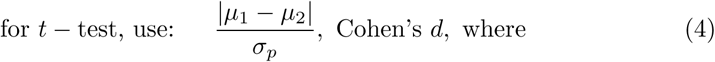

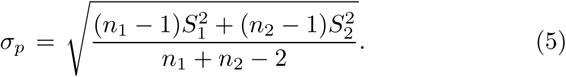

for Wilcoxon rank sum test, use: 
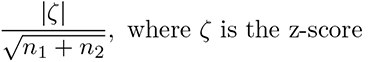

for one-way ANOVA, use: 
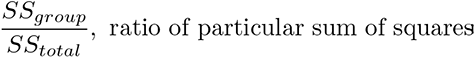

The variables *µ_j_* and 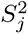 denote the sample mean and variances of the *j^th^* group. Effect sizes for *t* test and Wilcoxon rank sum test with values: (0, 0.2] are considered small, (0.2, 0.5] are medium, (0.5, 0.8] and above are large (Cohen, 2013; Tomczak and Tomczak, 2014). Effect sizes for one-way ANOVA with values (0, 0.01] are considered small, (0.01, 0.06] are medium, (0.06, 0.14] and above are large (Cohen, 2013).

## Declarations

### Data availability

Data is freely available at https://www.dx.doi.org/10.6084/m9.figshare.14877780.

### Code availability

Code is freely available at https://github.com/chengly70/SCDmodality.

### Author Contributions

WS, DS, and CL programmed and implemented models in software, including the data analysis. AB, WS, CL conceptualized and developed the project. SHG and WS designed the experiments and collected the data. DS and CL drafted the original manuscript. DS, AB, SHG, WS, CL edited the manuscript. CL supervised the project.

### Funding

This work was supported by the National Science Foundation (https://www.nsf.gov/): #IIS-1912338 DS and CL, #IIS-1912320 AB, #IIS-1912352 WS and SHG.

### Competing Interests

The authors declare that no competing interests exist. The funders had no role in study design, data collection and analysis, decision to publish, or preparation of the manuscript.

### Financial Interests

The authors declare they have no relevant financial or non-financial interests to disclose.

## Appendix A Projected correlation R_1_ theory

Here we consider how the CCA projected correlation between OB and aPC, which we denote by *R*_1_, is related to scaled error *E* (see Eqn (1)). These 2 entities should be negatively correlated because as *R*_1_ increases, the projected aPC activity should be ‘more like’ the OB activity, and the error should decrease as long as OB well-encodes modality and the noise correlation between OB and aPC is not too large. These practical considerations insure that *E* is not too large and is of limited range.

Similar to studying the co-variation between *E* and *ρ* in the main text, we only consider networks where: i) the OB (sub-)network well-encodes ortho vs. retro with optimal OB decoding accuracy ≥ 0.75; ii) the noise correlations are not too large: *ρ <* 0.15; iii) the standard deviation across all 8 *̺* values is *<* 0.2 for 2×2 (*<* 0.23 for 3×3). The resulting number of networks after these exclusions is large, see Table A1.

**Table A1.** Number of networks in projected correlation *R*_1_ theory shown in Figure A1.

|  | No Drug | Bicuculline | Muscimol |
| --- | --- | --- | --- |
| **Viable 2x2 for $R_1$ theory** | 80,455 | 63,437 | 26,619 |
| **Viable 3x3 for $R_1$ theory** | 3,292 | 2,503 | 1,581 |

Figure A1 shows scatter plots and normalized histograms that skew to the left, indicative of negative correlation between *E* and *R*_1_. The results are not as strong in the 3×3 networks compared to the 2×2 networks, but the summary statistics in Table A2 are pretty convincing. The results robustly hold across all three drug preparations.

**Table A2.**
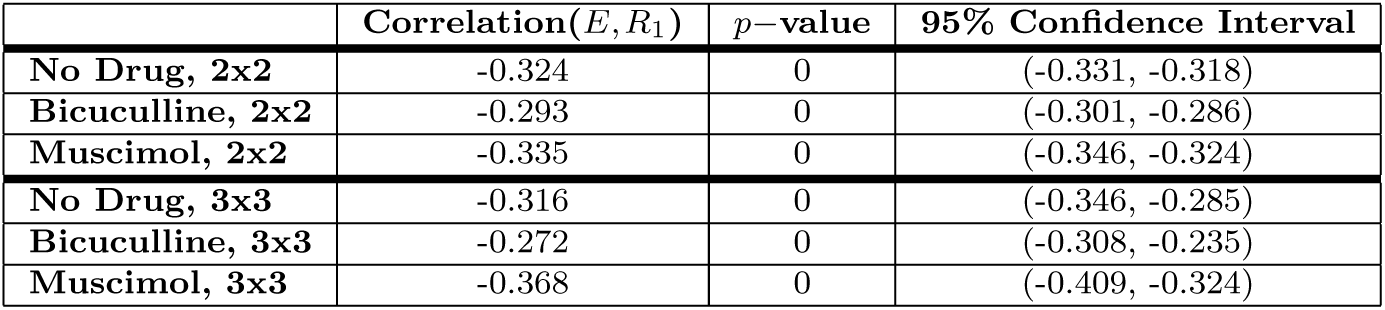
Statistics of (Pearson’s) correlation between *E* and CCA projected correlation *R*_1_. Our theory predicts that this relationship should be anti-correlated, which is evident by these statistics.

## Appendix B Population spike statistics

Next we show the pooled spike statistics over all neurons/pairs in aPC as well as cross-covariance/correlation between OB and aPC. The box plots in Figure B3 shows: i) comparisons between ortho and retro of the population mean, variance, covariance and (Pearson’s) correlation of spike counts across trials within the aPC region within a given drug preparation (A–D), ii) comparisons of drug preparation of spike count statistics within the aPC region for a given modality (I–L), iii) spike count cross-covariance and cross-correlation between OB and aPC comparisons of ortho versus retro with a given drug preparation (E–F), and comparisons of drug preparation within a given modality (G–H).

**Fig. A1.**
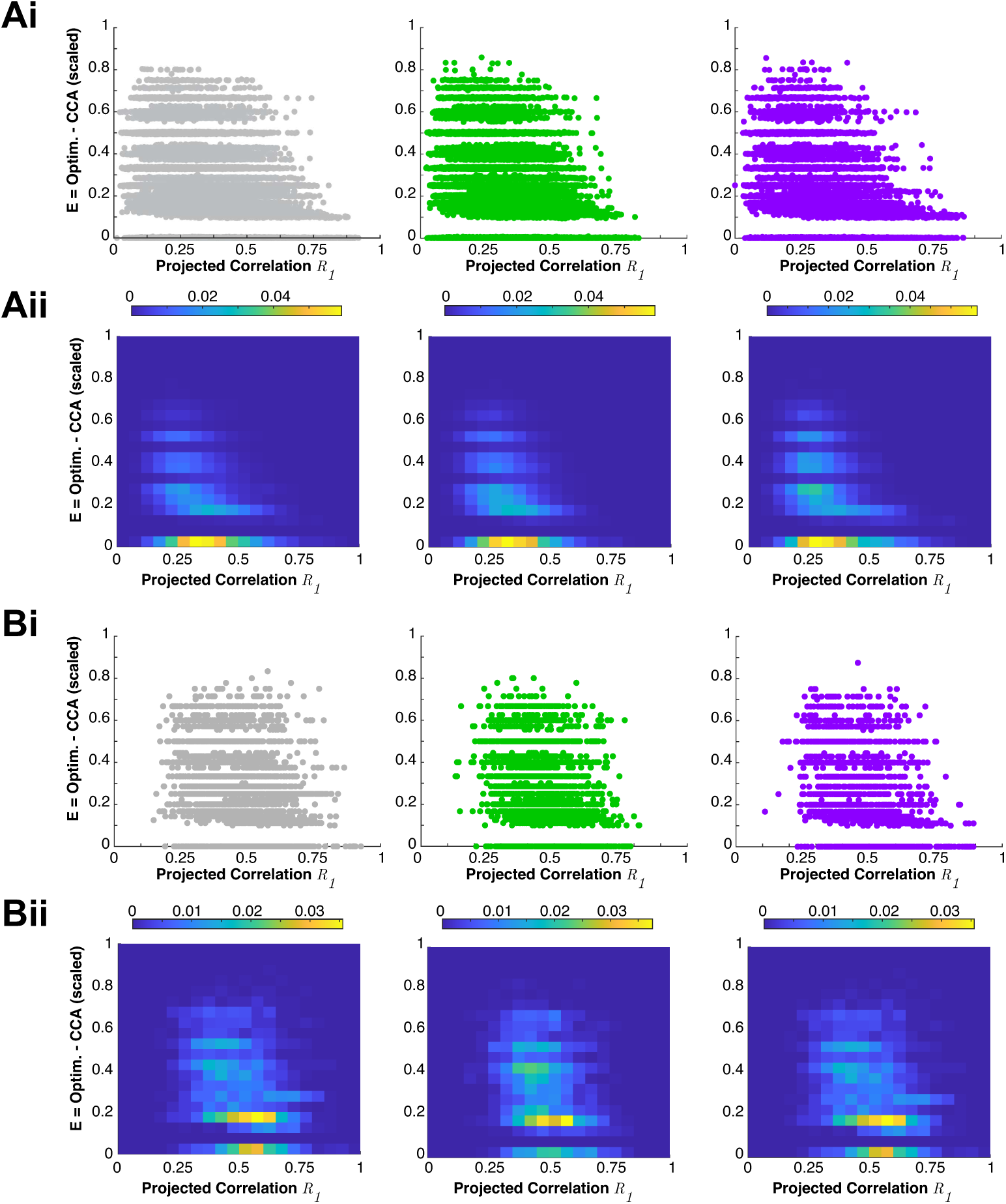
SCD error is negatively correlated with *R*_1_. A) For 2×2 networks, the error *E* (difference between optimal decoding accuracy and SCD) is negatively correlated with *R*_1_ (the CCA projected correlation between OB and aPC). Ai) shows a scatter plot of points (1 point for each network), Aii) shows the normalized histogram of (*R*_1_*, E*). See first row of Table A1 for counts and first 3 rows of Table A2 for summary statistics. B) Same as A) but for 3×3 networks; see second row of Table A1 for counts and last 3 rows of Table A2 for summary statistics.

**Fig. B2.**
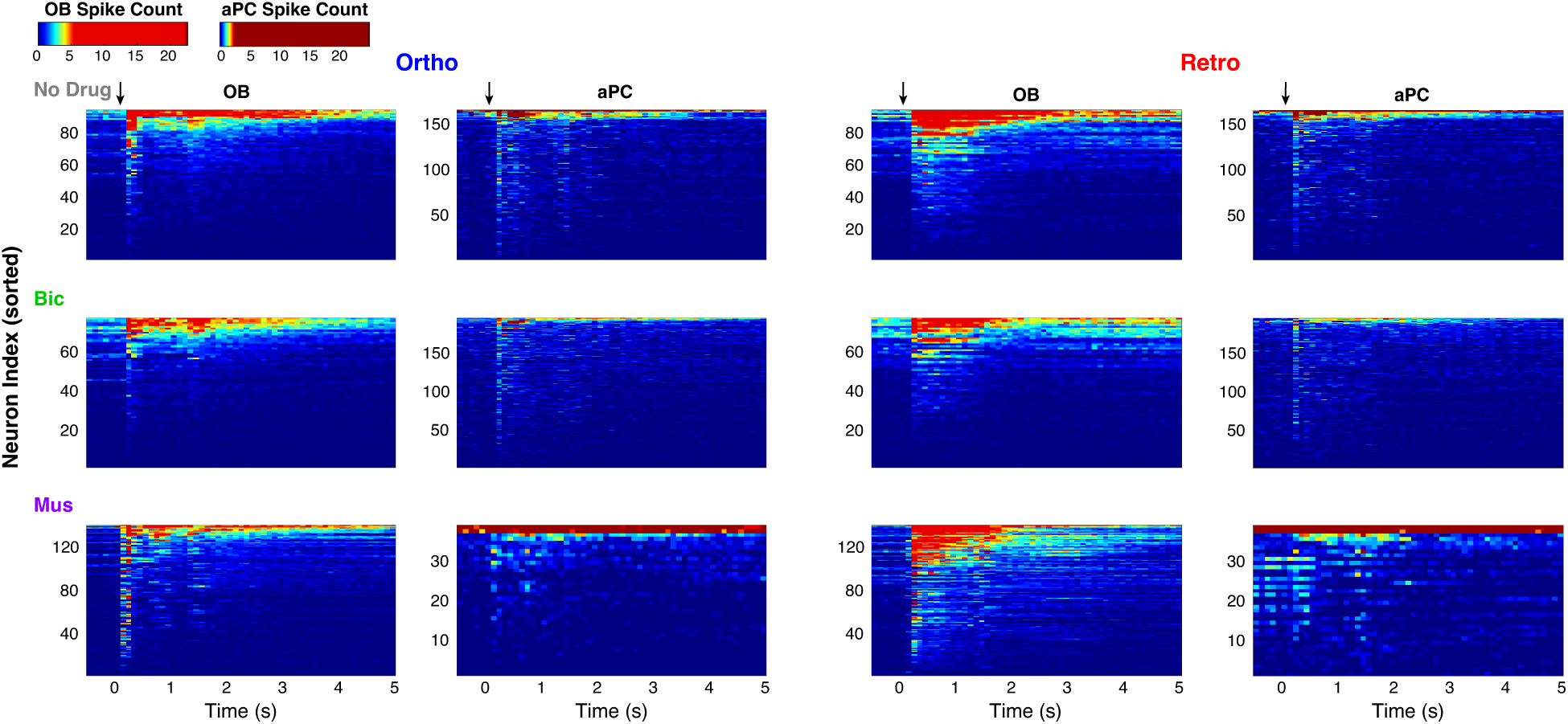
Representative population spiking in a rat. In a rat for each drug preparation, we show the spike count distribution in 100 ms windows as a function of time in both OB and aPC with both ortho and retro stimulation. For ND, there are 94 OB neurons and 165; for Bic there are 77 OB neurons and 195 aPC neurons; for Mus there are 140 OB neurons and 39 aPC neurons. The color bar legends are nonlinear for better contrast. Showing the entire set of recordings with all neurons (Table 1) or having the OB and aPC neurons on the same color bar scale would obscure the color-coding and be uninformative.

As stated in the Results section of this paper, with the large number of comparisons, the only differences that are statistically significant with reasonable (yet small) effect size are: spike count correlation with ortho stimulation comparing ND to Bic and Mus to Bic, spike count correlation with retro stimulation comparing ND to Bic (Fig B3L).

There are certainly instances where the population mean is vastly different (Fig B3B with Mus, C with Mus, J with retro, K with retro) but the long tail nature of the population statistics are likely the reason the effect sizes are minuscule.

The sparseness of the aPC responses measured by the fraction of neurons spiking in any given trial of odor presentation (Fig B4). Note that in a given recording, there are different numbers of simultaneously recorded neurons.

**Fig. B3.**
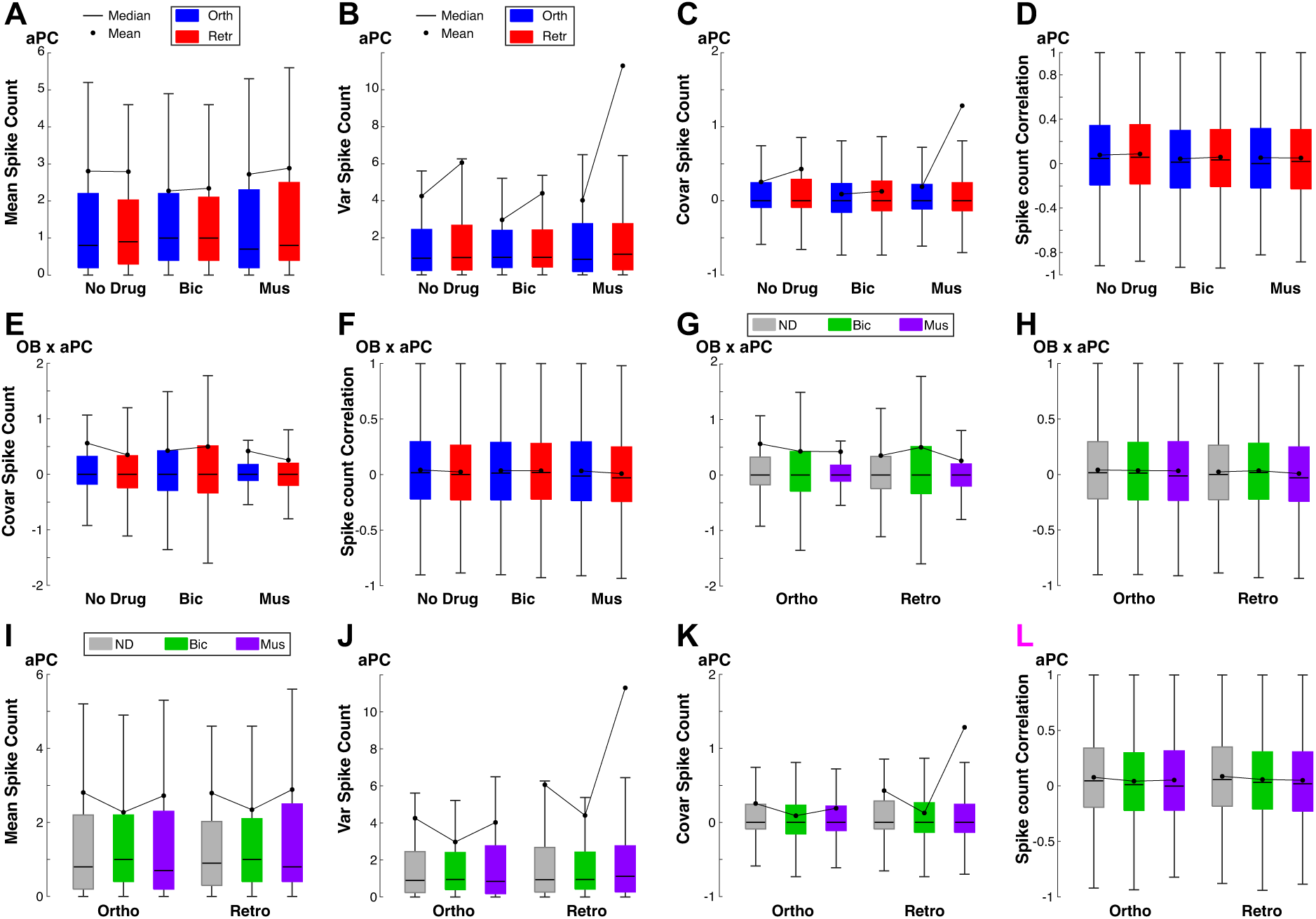
Population spike count statistics within aPC and for (OB,aPC). Box plots comparing population spike statistics: within aPC between modality within a given drug preparation (A–D), within aPC between drug preparations within a given modality (I–L), cross-covariance and correlation between OB and aPC comparing modality within a given drug preparation (E–F), and comparing drug preparation within a modality (G–H). See Tables 1 and 2 (first three rows) for number of individual neurons and pairs.

## Appendix C Detailed Statistics of Results

Here we show the details for the three statistical tests: *p*−values and effect sizes.

**Fig. B4.**
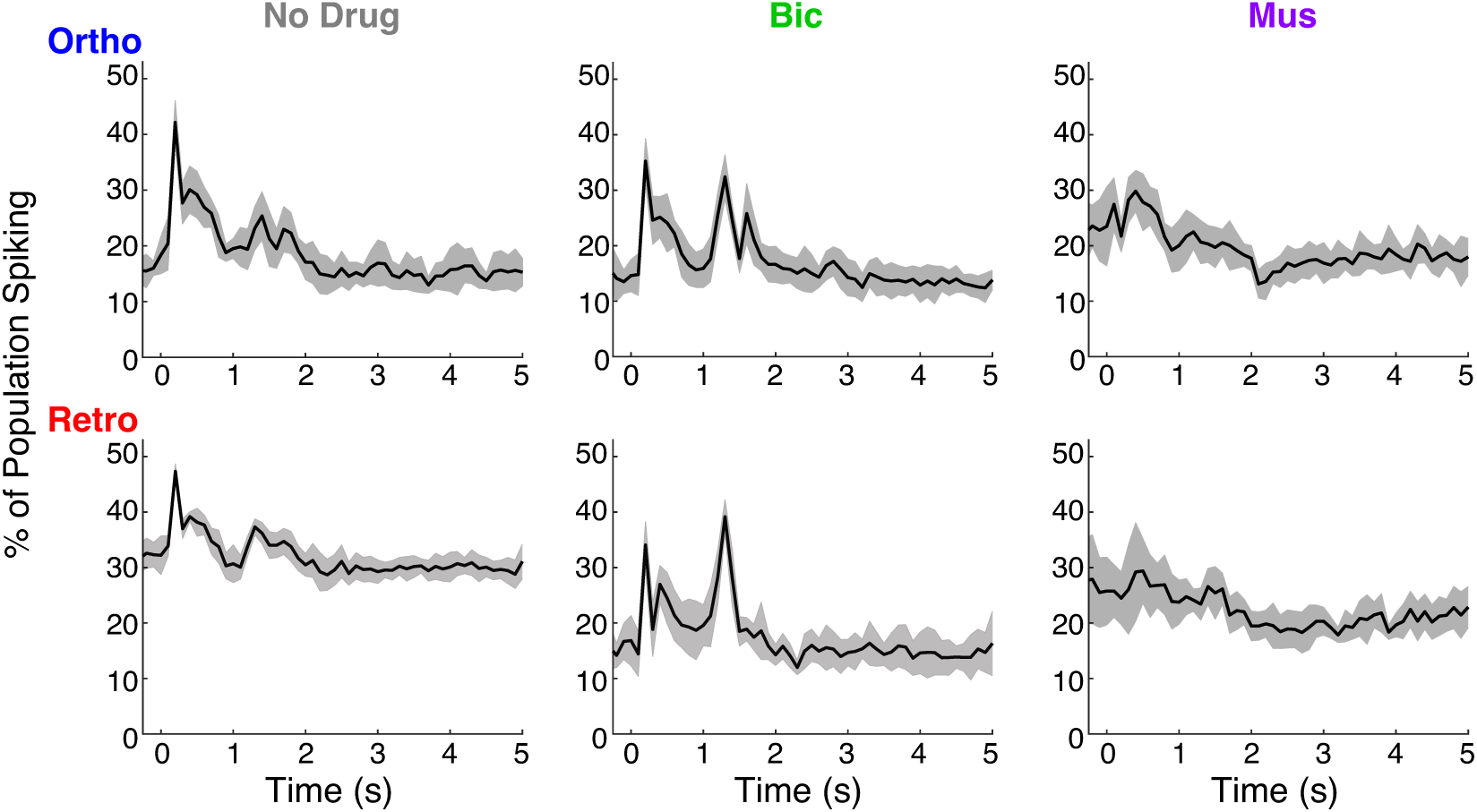
The sparseness of aPC responses.

Showing the fraction of aPC neurons spiking in a given trial as a function of time. The black curve is the average over all trials, and shaded region is a measure of the standard deviation across all trials.

**Table C3.** Statistic to show that OB has better single neuron decoding accuracy than aPC; see Figure 1C. Using 3 tests: two-sample *t*−test assuming unequal variance, Wilcoxon rank sum test, and one-way ANOVA. Bottom table shows effect size.

| Relationship / $p$-value | $t$-test | Rank Sum | ANOVA |
| --- | --- | --- | --- |
| ND | $1.7 \times 10^{-47}$ | $6.8 \times 10^{-40}$ | $1.3 \times 10^{-47}$ |
| Bic | $1.5 \times 10^{-12}$ | $9.2 \times 10^{-12}$ | $5.2 \times 10^{-13}$ |
| Mus | $3.5 \times 10^{-12}$ | $1.6 \times 10^{-5}$ | $4.5 \times 10^{-10}$ |
| Relationship / Effect Size | $t$-test | Rank Sum | ANOVA |
| ND | 0.70 (large) | 0.31 (medium) | 0.11 (large) |
| Bic | 0.49 (medium) | 0.23 (medium) | 0.06 (small/medium) |
| Mus | 0.44 (medium) | 0.15 (small) | 0.05 (small) |

**Table C4.** Statistic to show that CCA is better than LDA and that optimal is better than CCA for 2×2 networks with 1 s time window; see Figures 2C and 3A. Using 3 tests: two-sample *t*−test assuming unequal variance, Wilcoxon rank sum test, and one-way ANOVA. We only show the effect size because the *p*−values are 0 for both comparisons with machine precision using double.

| Relationship / Effect Size | $t$-test | Rank Sum | ANOVA |
| --- | --- | --- | --- |
| ND: LDA<CCA | 0.59 (large) | 0.04 (small) | 0.001 (small) |
| Bic: LDA<CCA | 0.88 (large) | 0.05 (small) | 0.003 (small) |
| Mus: LDA<CCA | 0.72 (large) | 0.04 (small) | 0.002 (small) |
| Relationship / Effect Size | $t$-test | Rank Sum | ANOVA |
| ND: CCA<Optim | 0.87 (large) | 0.05 (small) | 0.003 (small) |
| Bic: CCA<Optim | 0.86 (large) | 0.05 (small) | 0.003 (small) |
| Mus: CCA<Optim | 0.85 (large) | 0.05 (small) | 0.003 (small) |

**Table C5.** Statistic to show that CCA is better than LDA and that optimal is better than CCA for 3×3 networks with 1 s time window; see Figure 4A. Using 3 tests: two-sample *t*−test assuming unequal variance, Wilcoxon rank sum test, and one-way ANOVA. We only show the effect size because the *p*−values are numerically 0 for both comparisons with machine precision using double.

| Relationship / Effect Size | $t$-test | Rank Sum | ANOVA |
| --- | --- | --- | --- |
| ND: LDA<CCA | 0.50 (large) | 0.18 (small) | 0.03 (medium) |
| Bic: LDA<CCA | 0.80 (large) | 0.28 (medium) | 0.08 (large) |
| Mus: LDA<CCA | 0.61 (large) | 0.13 (small) | 0.02 (medium) |
| Relationship / Effect Size | $t$-test | Rank Sum | ANOVA |
| ND: CCA<Optim | 1.40 (large) | 0.43 (medium) | 0.20 (large) |
| Bic: CCA<Optim | 1.39 (large) | 0.45 (medium) | 0.21 (large) |
| Mus: CCA<Optim | 1.32 (large) | 0.28 (medium) | 0.09 (large) |

## Footnotes

1 The condition iii) was changed to include networks with standard deviation < 0.23, taken across 18 values compared to 8 values in 2x2 networks; this insured we had at least 1000 viable 3x3 networks.

## Notes

### Competing Interest Statement

The authors have declared no competing interest.

### Summary of Updates

Added a new figure (6), extra appendix and corresponding tables, changes in text, etc.

